# Augmentation of CD47-SIRPα signaling protects cones in genetic models of retinal degeneration

**DOI:** 10.1101/2021.04.23.440841

**Authors:** Sean K. Wang, Yunlu Xue, Constance L. Cepko

## Abstract

Inherited retinal diseases such as retinitis pigmentosa (RP) can be caused by thousands of different mutations, a small number of which have been successfully treated with gene replacement. However, this approach has yet to scale and may be infeasible in many cases, highlighting the need for interventions that could benefit more patients. Here, we found that microglial phagocytosis is upregulated during cone degeneration in RP, suggesting that expression of “don’t eat me” signals such as CD47 might confer protection to cones. To test this, we delivered an adeno-associated viral (AAV) vector expressing CD47 on cones, which promoted cone survival in three mouse models of RP and preserved visual function. Cone rescue with CD47 required signal regulatory protein alpha (SIRPα) but not microglia or thrombospondin-1 (TSP1), suggesting that CD47 interacts with SIRPα on non-microglial cells to alleviate degeneration. These findings establish augmentation of CD47-SIRPα signaling as a potential treatment strategy for RP and possibly other forms of neurodegeneration.

## INTRODUCTION

A major obstacle in developing treatments for inherited retinal diseases (IRDs) is the enormous genetic heterogeneity of pathogenic mutations. In the largest family of these disorders, retinitis pigmentosa (RP), thousands of unique mutations have been identified spanning ∼100 different genes (https://sph.uth.edu/retnet/). For a minority of IRDs, gene replacement therapy with adeno-associated viral (AAV) vectors has led to promising results in clinical trials, and in one instance, a successful commercial drug (1, 2). Nonetheless, this strategy has yet to reach the vast majority of patients and may not be feasible for those with less common, gain-of-function, or unidentified mutations.

Given these challenges, another approach to treating RP and other IRDs could be to develop mutation-agnostic therapies that preserve cone photoreceptors. In RP, disease mutations, many of which are exclusively expressed in rods, lead to the early death of these cells, often before the condition is diagnosed (3). Rod death is in itself tolerable, as rods are only needed for vision in dim light. However, rod loss is followed by the degeneration of cones, resulting in deterioration of daylight, color, and high-acuity vision. The reasons for secondary cone degeneration in RP remain incompletely understood, but likely include increased oxidative stress (4, 5), decreased glucose availability (6–8), and immune dysregulation (9, 10). Regardless of the exact etiology, an intervention capable of protecting cones in RP independent of the initial genetic lesion would potentially be beneficial for many patients.

One of the hallmarks of neurodegenerative disorders including RP is the activation of microglia, the resident immune cells of the retina and central nervous system (11, 12). This activation upregulates phagocytosis and inflammatory cytokine production, facilitating the clearance of infections and cell debris, but can also damage nearby healthy tissue. In mice, activated microglia during the early stages of RP can engulf living rods, thereby hastening rod death (13, 14). While microglia are similarly activated during cone degeneration (9, 10), their impact on cone survival appears to be more nuanced. Previously, we showed that the anti-inflammatory cytokine TGF-β1 requires microglia to preserve cones in RP mouse models (10), indicating that microglia are capable of conferring protection to cones. However, we also found the net effect of microglia depletion on cone degeneration to be minimal (9, 10), suggesting that any beneficial activities of microglia on cones are likely counter-balanced by detrimental ones. These findings left open the role of microglia in cone death and led us to ask whether microglial phagocytosis might contribute to cone demise.

Here, we tested if expression of an anti-phagocytic molecule on cones could slow the rate of cone degeneration. Specifically, we used an AAV vector encoding CD47, a “don’t eat me” signal known to inhibit engulfment via interaction with signal regulatory protein alpha (SIRPα) on phagocytic cells (15, 16). Expression of CD47 preserved cones and vision in multiple mouse models of RP through a mechanism dependent on SIRPα, but surprisingly not microglia. Our findings support augmentation of CD47-SIRPα signaling as a possible mutation-agnostic therapy for RP and potentially other neurodegenerative diseases.

## RESULTS

### Microglia show increased phagocytic activity during secondary cone degeneration

We previously generated *rd1*;CX3CR1^GFP/+^ and sighted CX3CR1^GFP/+^ (*rd1* heterozygous) mice by crossing the widely used *rd1* model of RP with CX3CR1^GFP^ animals, in which microglia express GFP (9, 17, 18). To assess if microglia during the later stages of RP might phagocytose cones, retinas from these mice were immunostained for cone arrestin, a marker of all cones. In sighted CX3CR1^GFP/+^ retinas, microglia lacked any appreciable contact with cone arrestin-positive cells (Figure 1A), consistent with the normal exclusion of microglia from the photoreceptor layer (19). In contrast, microglia in *rd1*;CX3CR1^GFP/+^ retinas were often directly adjacent to degenerating cones, although in no case was overt engulfment of cones observed. To more sensitively measure microglial phagocytic activity, we explanted retinas from *rd1*;CX3CR1^GFP/+^ and sighted CX3CR1^GFP/+^ animals and incubated them with yeast particles conjugated to pHrodo Red, a pH-dependent dye that fluoresces upon lysosomal acidification (Figure 1B) (20). Microglia from these retinas were then analyzed by flow cytometry at postnatal day 20 (P20), the approximate age at which cone death in *rd1* mice begins (6), as well as P50, after substantial cone loss has occurred. At both time points, microglia from *rd1*;CX3CR1^GFP/+^ retinas internalized significantly more yeast than those from sighted controls (Figure 1C), suggesting persistent elevation of microglial uptake during the later stages of RP. Microglia thus exhibit increased phagocytic activity during secondary cone degeneration, a factor we hypothesized might worsen cone demise.

**Figure 1.**
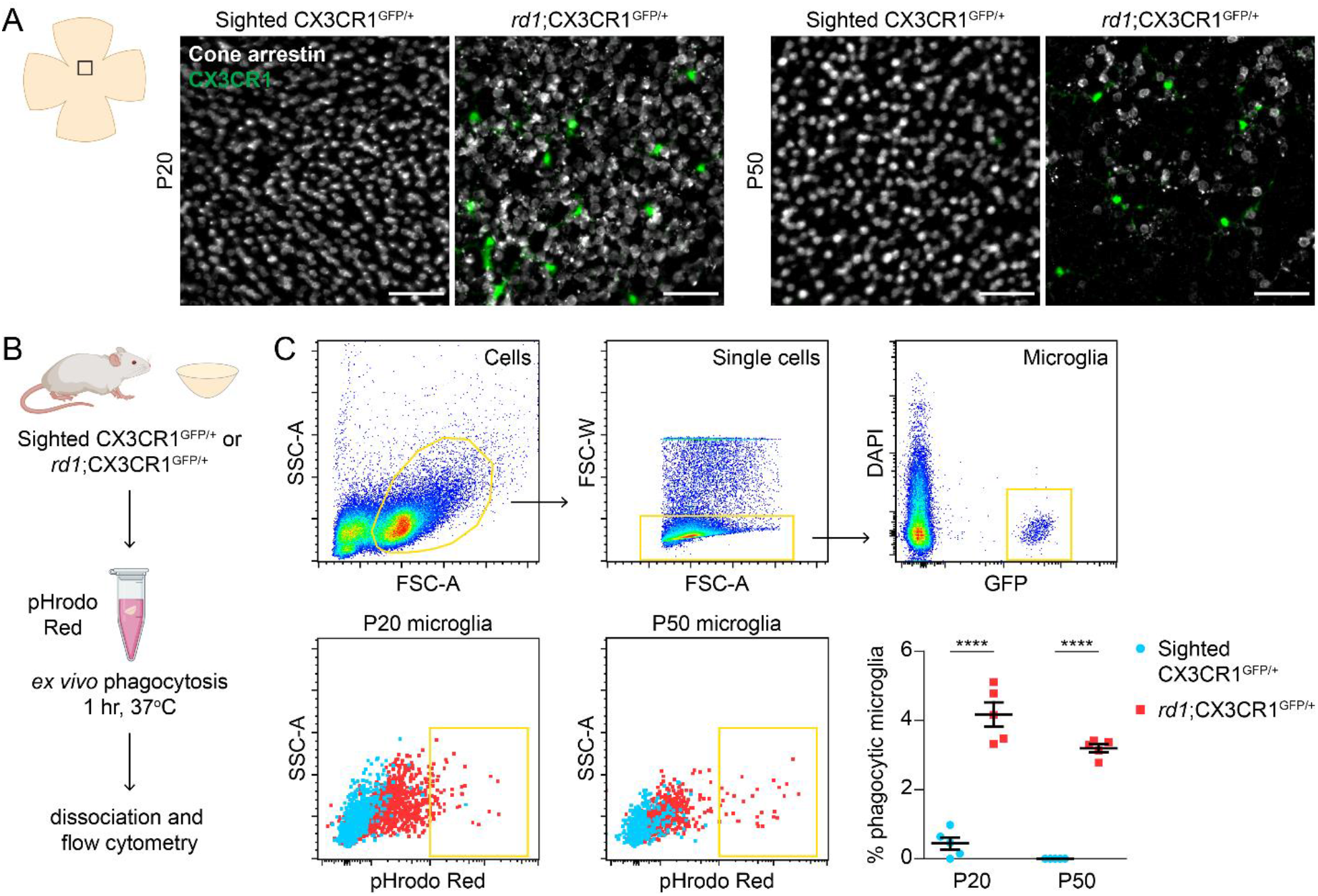
Cone degeneration is associated with increased microglial phagocytosis. (**A**) Cone arrestin immunostaining and CX3CR1-positive microglia in flat-mounted retinas from sighted CX3CR1^GFP/+^ (*rd1* heterozygous) and *rd1*;CX3CR1^GFP/+^ mice at P20 and P50. Images were acquired from the central retina. Scale bars, 50 µm. (**B**) Schematic of *ex vivo* phagocytosis assay. Retinas from sighted CX3CR1^GFP/+^ and *rd1*;CX3CR1^GFP/+^ mice were incubated with yeast (zymosan) particles conjugated to pHrodo Red, a pH-sensitive dye that fluoresces upon lysosomal acidification. Microglia were subsequently analyzed by flow cytometry. (**C**) Flow cytometry gating for microglia and quantification of microglial phagocytosis in sighted CX3CR1^GFP/+^ (*n* = 5) and *rd1*;CX3CR1^GFP/+^ (*n* = 5) retinas at P20 and P50. Data are shown as mean ± SEM. **** *P*<0.0001 by two-tailed Student’s t-test.

### Expression of the CD47 “don’t eat me” signal promotes survival of degenerating cones

Among the key regulators of phagocytosis are “don’t eat me” signals such as CD47, which when present on cells, impede their engulfment by macrophages (21). To test if inhibiting phagocytosis during RP might benefit cones, we created an AAV vector (AAV8-RedO-CD47) using the human red opsin promoter to express CD47 on cones (Figure 2A). In wild-type mice injected subretinally with a GFP control vector (AAV8-RedO-GFP), which labels both M- and S-type cones (8), endogenous CD47 could be seen in retinal plexiform layers as previously described, but not in photoreceptors (Figure 2B) (22). Following co-administration of AAV8-RedO-GFP plus AAV8-RedO-CD47, CD47 immunostaining could additionally be detected in cones. Notably, use of AAV8-RedO-CD47 in wild-type mice produced no obvious changes in cone or retinal morphology when evaluated one month later.

**Figure 2.**
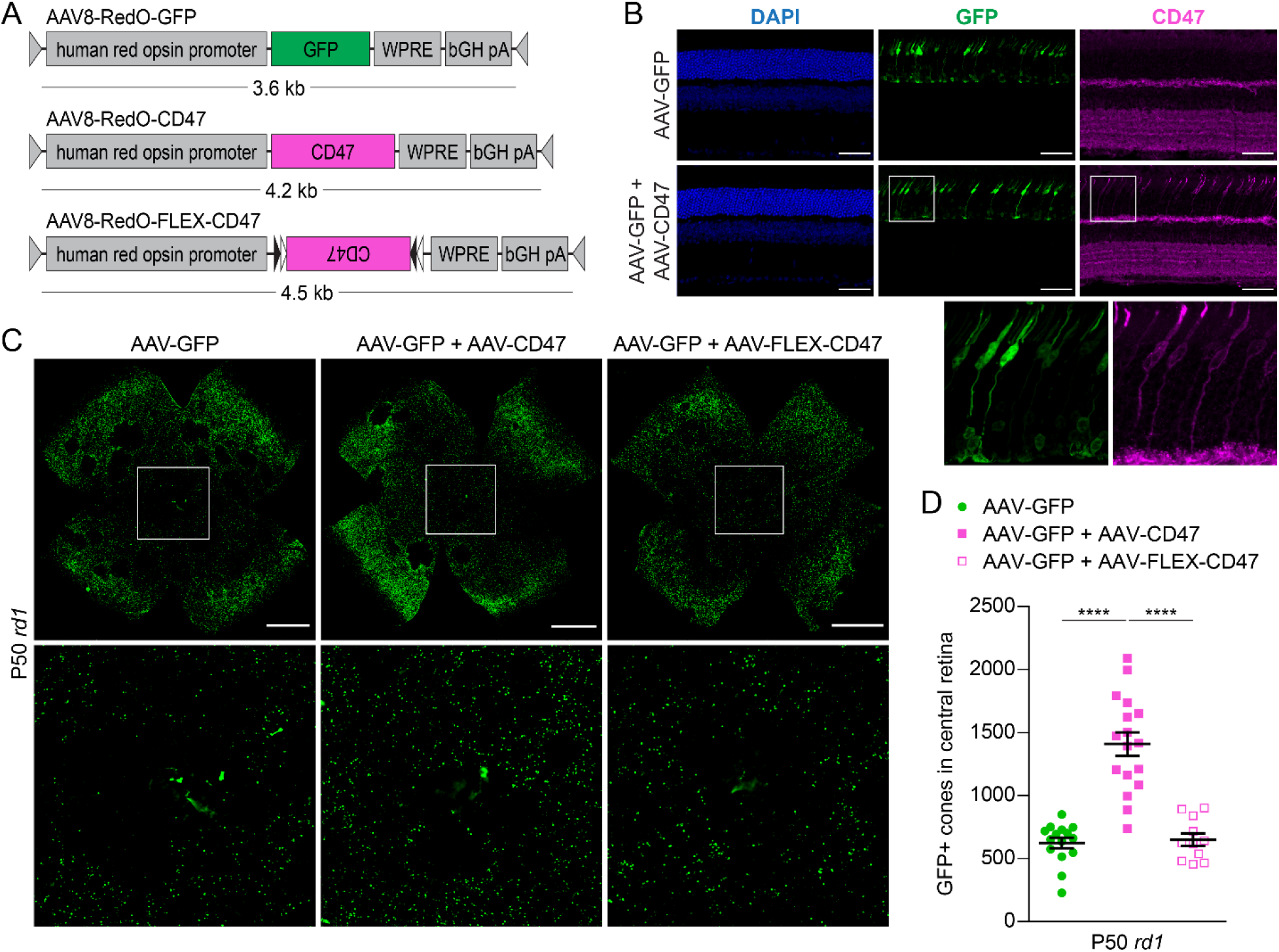
Effect of CD47 expression on cone survival. (**A**) Schematics of AAV vectors. AAV8-RedO-FLEX-CD47 is a flip-excision (FLEX) vector in which the CD47 transgene is inverted and flanked by lox2272 (black triangles) and loxP (white triangles) sites. (**B**) Immunostaining for CD47 in P40 wild-type (CD-1) retinas following infection with AAV8-RedO-GFP or AAV8-RedO-GFP plus AAV8-RedO-CD47. Nuclei were labeled with 4′,6-diamidino-2-phenylindole (DAPI). Scale bars, 50 µm. (**C**) Representative flat-mounts of P50 *rd1* retinas following infection with AAV8-RedO-GFP, AAV8-RedO-GFP plus AAV8-RedO-CD47, or AAV8-RedO-GFP plus AAV8-RedO-FLEX-CD47. Paired images depict low and high magnifications. Scale bars, 1 mm. (**D**) Quantification of GFP-positive cones in central retinas of *rd1* mice (*n* = 11-17) following infection with AAV8-RedO-GFP, AAV8-RedO-GFP plus AAV8-RedO-CD47, or AAV8-RedO-GFP plus AAV8-FLEX-RedO-CD47. Data are shown as mean ± SEM. **** *P*<0.0001 by two-tailed Student’s t-test.

In mouse models of RP, cone degeneration proceeds from the optic nerve head outward with relative sparing of the peripheral retina. To measure the effect of CD47 on cone survival, GFP-positive cones in the central retinas of *rd1* mice were quantified. Cone counting using a GFP vector has been previously validated by our group and provides unambiguous labeling of cones, enabling automated quantification (4, 9, 10, 23). At these doses, co-infection with two serotype 8 vectors in cones is expected to be at least 90% (4). Compared to AAV8-RedO-GFP alone, co-infection with AAV8-RedO-CD47 approximately doubled the number of cones in the central retina at P50 (Figure 2, C and D). In contrast, co-infection with AAV8-RedO-FLEX-CD47, a control vector with the CD47 sequence inverted, did not significantly change the number of remaining cones. To assess cone preservation with CD47 beyond the central retina, entire *rd1* retinas were next analyzed for GFP-positive cones using flow cytometry (Supplemental Figure 1). Consistent with the histological findings, this method showed greater cone counts at P50 with AAV8-RedO-GFP plus AAV8-RedO-CD47 than AAV8-RedO-GFP only (Supplemental Figure 1). To also gauge cone survival versus untreated eyes, *rd1* retinas at P50 were immunostained for cone arrestin. Relative to central retinas without treatment or receiving AAV8-RedO-GFP alone, addition of AAV8-RedO-CD47 again resulted in more cones as defined by this marker (Supplemental Figure 2). Together, these data demonstrated that CD47 expression could promote survival of cones in *rd1* mice.

### CD47 promotes cone survival and retention of vision in multiple genetic models

To determine if CD47 might similarly slow retinal degeneration in other models of RP, AAV8-RedO-CD47 was tested in *rd10* mice, which carry a missense mutation in *Pde6b*, and *Rho*^*-/-*^ mice, which lack rhodopsin. In both *rd10* retinas at P100 and P130 and *Rho*^*-/-*^ retinas at P150, AAV8-RedO-CD47 again improved the number of cones (Figure 3, A-C and Supplemental Figure 1), suggesting that CD47 may generically combat cone degeneration. To investigate the therapeutic relevance of AAV8-RedO-CD47, treated animals were then subjected to two visually dependent behavioral assays. First, a light-dark discrimination test was performed by leveraging the natural preference of sighted mice for dark rather than well-illuminated spaces (see Methods). In keeping with this, wild-type animals spent ∼70% of their time in the dark half of a 50:50 light-dark environment, while *rd1* mice without treatment or receiving only AAV8-RedO-GFP divided their time evenly between the two chambers (Figure 3D). Compared to these latter two groups, *rd1* animals receiving AAV8-RedO-GFP plus AAV8-RedO-CD47 spent significantly more time in the dark, consistent with better preservation of visual function. As a second measure of vision, mice were evaluated using an optomotor assay in which moving stripes were presented to elicit the visually dependent optomotor response. By varying the spatial frequency of stripes to make them easier or more difficult to see, the visual threshold in each eye could be estimated (24). In *rd10* animals tested at P60, visual thresholds were significantly higher in eyes infected with AAV8-RedO-GFP plus AAV8-RedO-CD47 than AAV8-RedO-GFP alone (Figure 3E), again suggesting better retention of sight. CD47 expression thus not only promotes cone survival in different mouse models of RP, but also protects from vision loss, supporting its potential use as a mutation-agnostic therapy for this condition.

**Figure 3.**
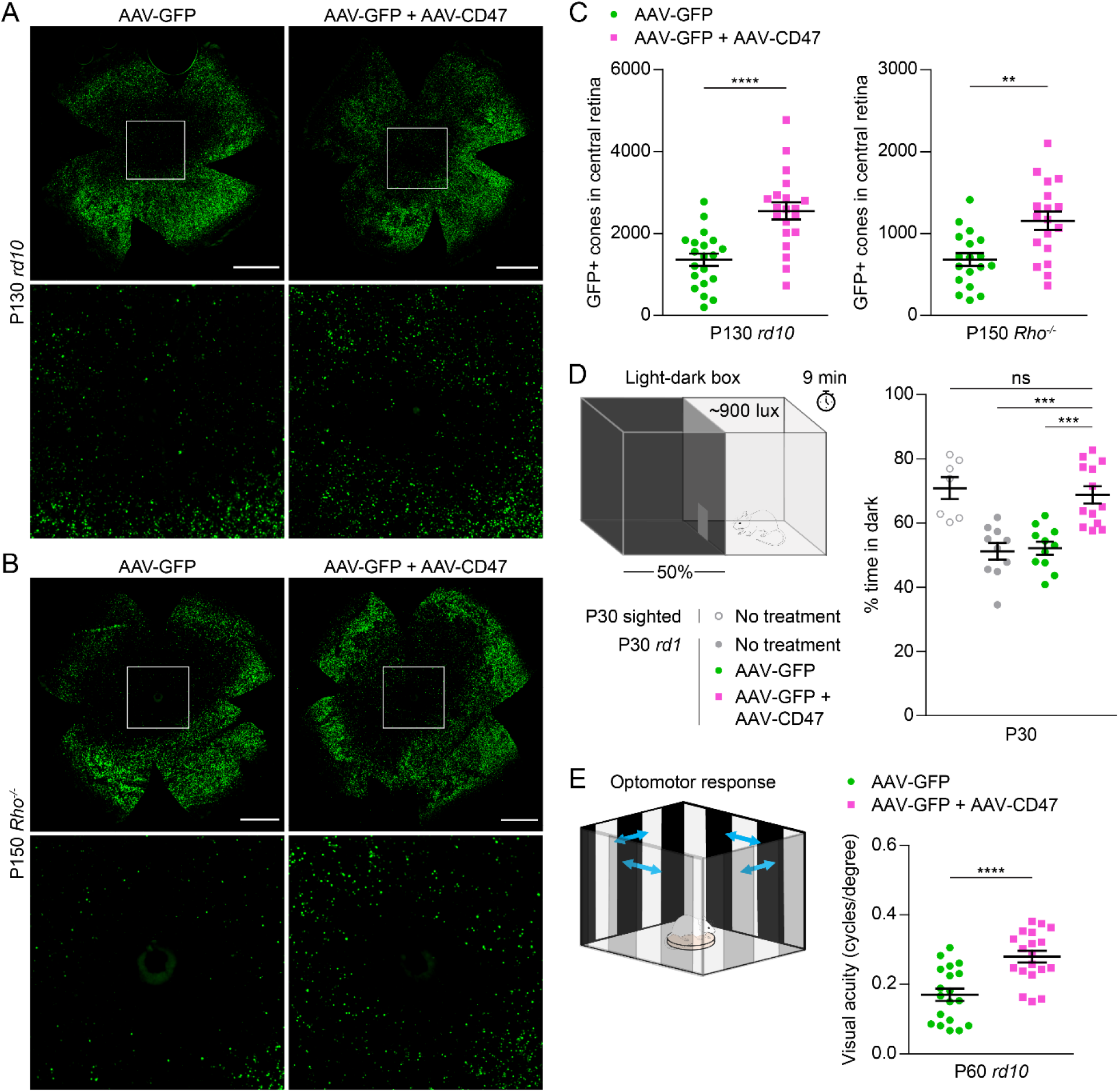
Effect of CD47 expression on long-term cone survival and visual function. (**A**) Representative flat-mounts of P130 *rd10* retinas following infection with AAV8-RedO-GFP or AAV8-RedO-GFP plus AAV8-RedO-CD47. Paired images depict low and high magnifications. Scale bars, 1 mm. (**B**) Representative flat-mounts of P150 *Rho*^*-/-*^ retinas following infection with AAV8-RedO-GFP or AAV8-RedO-GFP plus AAV8-RedO-CD47. Paired images depict low and high magnifications. Scale bars, 1 mm. (**C**) Quantification of GFP-positive cones in central retinas of *rd10* (*n* = 20) and *Rho*^*-/-*^ (*n* = 18) mice following infection with AAV8-RedO-GFP or AAV8-RedO-GFP plus AAV8-RedO-CD47. (**D**) Percent time spent in dark in a 50:50 light-dark box for untreated (*n* = 7-10) and *rd1* (*n* = 11-13) mice following infection with AAV8-RedO-GFP or AAV8-RedO-GFP plus AAV8-RedO-CD47. (**E**) Visual thresholds in eyes from P60 *rd10* mice (*n* = 19) as measured by optomotor following infection with AAV8-RedO-GFP or AAV8-RedO-GFP plus AAV8-RedO-CD47. Data are shown as mean ± SEM. ** *P*<0.01, *** *P*<0.001, **** *P*<0.0001 by two-tailed Student’s t-test for (C) and (E), two-tailed Student’s t-test with Bonferroni correction for (D). ns, not significant.

### Delayed expression of CD47 ameliorates cone death

Clinically, patients with RP are often diagnosed following the development of night blindness (3). We therefore asked whether CD47 could still protect cones even after the majority of rods have died. To model this scenario, we bred R26-CreERT2 mice, which undergo inducible Cre activation in the presence of tamoxifen, with the *rd1* strain to obtain *rd1*;CreERT2/+ animals. When combined with a flip-excision (FLEX) vector (25), this approach allowed for delayed AAV expression while avoiding the technical challenges of subretinal injections in older mice. As a test for CreERT2-mediated recombination, P0-1 *rd1*;CreERT2/+ animals were infected with AAV8-RedO-FLEX-mCherry, a vector designed to express mCherry only after exposure to tamoxifen (Figure 4A). In the absence of tamoxifen, AAV8-RedO-FLEX-mCherry produced no detectable fluorescence after 30 days (Figure 4B). However, the same vector with tamoxifen from P19-21 led to robust mCherry expression throughout the retina. Using this strategy and an analogous AAV8-RedO-FLEX-CD47 vector, the effect of delayed CD47 expression on cone survival was examined. In *rd1*;CreERT2/+ animals co-infected with AAV8-RedO-GFP plus AAV8-RedO-FLEX-CD47 but without tamoxifen, the number of cones at P50 was comparable to that seen in *rd1* mice (Figure 2D and Figure 4, C and D). In contrast, in animals additionally receiving tamoxifen from P19-21, an age by which most rods have died (6), significantly more GFP-positive cones were observed. Collectively, these findings suggest that CD47 may alleviate cone degeneration even if administered later in the disease progression.

**Figure 4.**
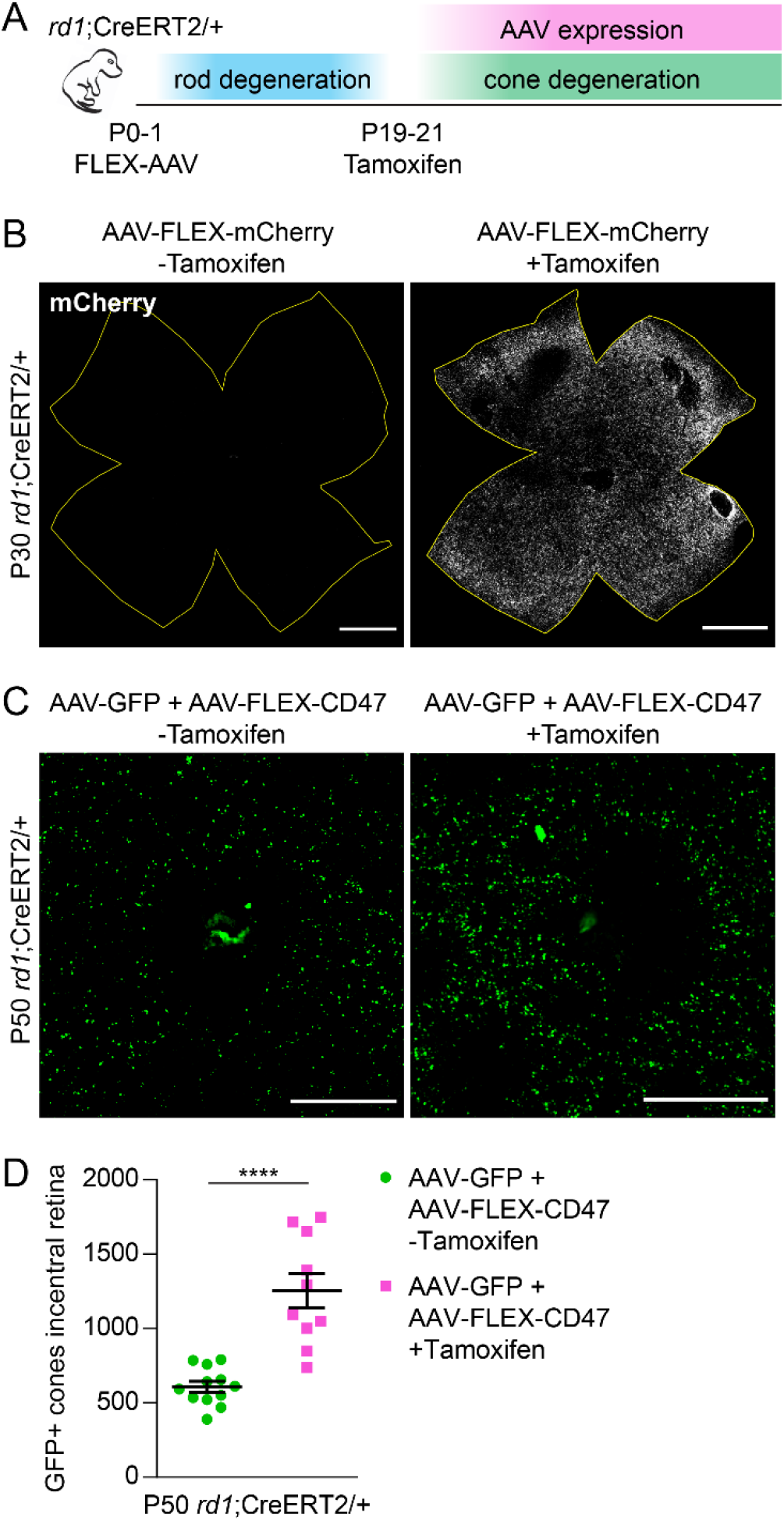
Effect of delayed CD47 expression on cone survival. (**A**) Schematic of delayed AAV expression experiments. P0-1 *rd1*;CreERT2/+ mice were subretinally injected with FLEX vectors, which were subsequently activated by intraperitoneal (IP) injections of tamoxifen from P19-21. (**B**) Representative flat-mounts of P30 *rd1*;CreERT2/+ retinas following infection with AAV8-RedO-FLEX-mCherry with or without IP tamoxifen. Boundaries of each retina are depicted in yellow. Scale bars, 1 mm. (**C**) Representative images of central retinas from P50 *rd1*;CreERT2/+ mice following infection with AAV8-RedO-GFP plus AAV8-RedO-FLEX-CD47 with or without IP tamoxifen. Scale bars, 500 µm. (**D**) Quantification of GFP-positive cones in central retinas of *rd1*;CreERT2/+ mice (*n* = 10-12) following infection with AAV8-RedO-GFP plus AAV8-RedO-FLEX-CD47 with or without IP tamoxifen. Data are shown as mean ± SEM. **** *P*<0.0001 by two-tailed Student’s t-test.

### CD47 requires SIRPα but not microglia or TSP1 to preserve cones

In the brain, the “don’t eat me” function of CD47 is thought to be largely mediated by microglia (16, 26). We thus suspected that AAV8-RedO-CD47 might protect cones by inhibiting their phagocytosis by microglia. To investigate this mechanism, mice infected subretinally with AAV8-RedO-CD47 had their microglia pharmacologically depleted using PLX5622, a small molecule blocking a receptor on microglia essential for their survival (27). In the retina, PLX5622 depleted ∼99% of microglia within 15 days (Figure 5, A and B), demonstrating virtually complete elimination of this cell type. Consistent with our previous observations (10), PLX5622 administration in *rd1* mice from P20-49 had no significant effect on cone counts in eyes receiving only AAV8-RedO-GFP (Figure 5, C and D). Surprisingly, PLX5622 also failed to perturb cone preservation with AAV8-RedO-CD47, indicating that microglia were dispensable for its mechanism of action.

**Figure 5.**
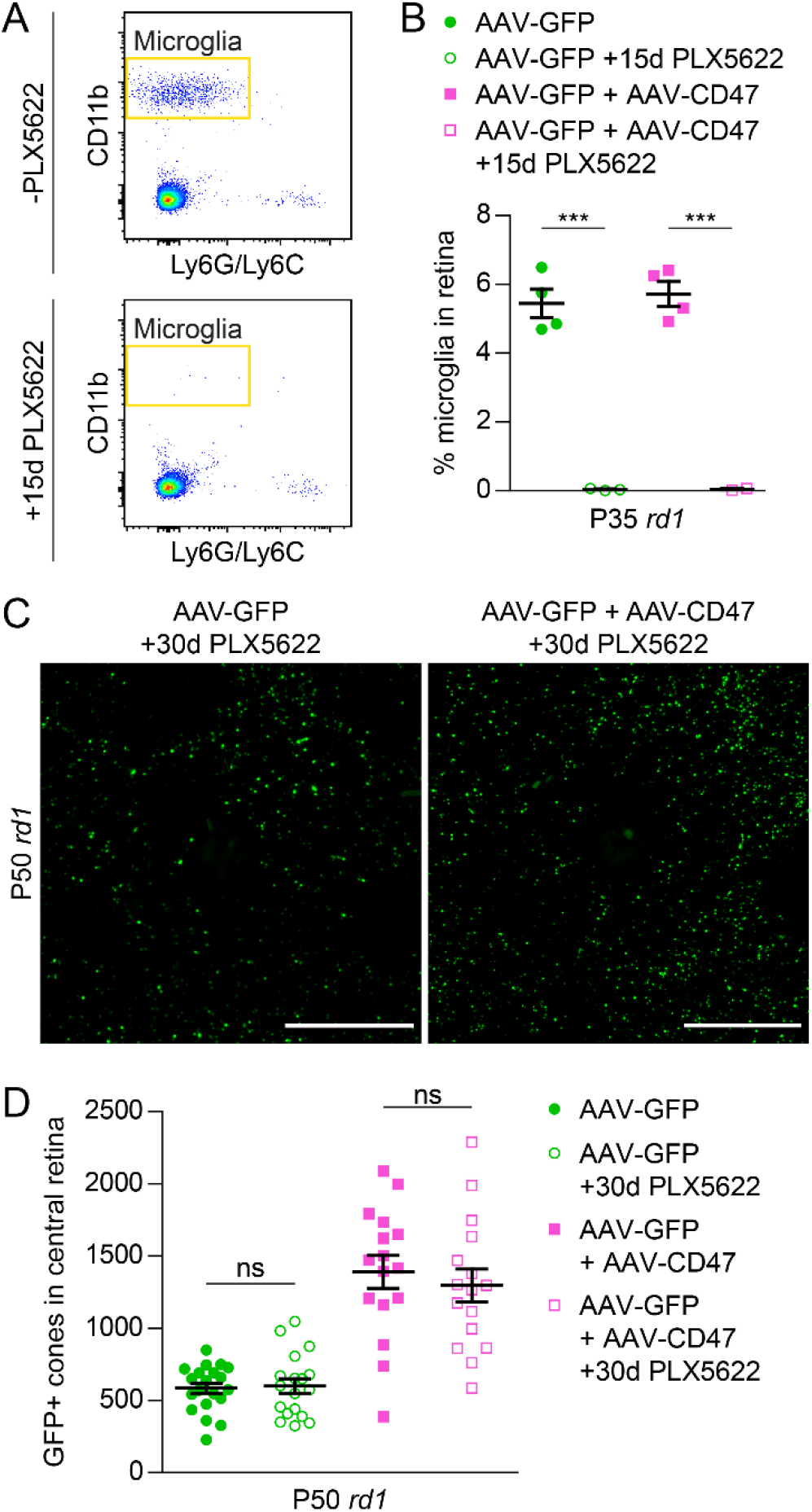
Role of microglia in CD47-mediated cone survival. (**A**) Flow cytometry gating for retinal microglia from P35 *rd1* mice with or without PLX5622 from P20-34. Microglia were defined as CD11b-positive Ly6G/Ly6C-negative cells. (**B**) Quantification by flow cytometry of retinal microglia from P35 *rd1* mice (*n* = 2-4) following infection with AAV8-RedO-GFP or AAV8-RedO-GFP plus AAV8-RedO-CD47 with or without PLX5622 from P20-34. (**C**) Representative images of central retinas from P50 *rd1* mice following infection with AAV8-RedO-GFP or AAV8-RedO-GFP plus AAV8-RedO-CD47 and PLX5622 from P20-49. Scale bars, 500 µm. (**D**) Quantification of GFP-positive cones in central retinas of *rd1* mice (*n* = 16-18) following infection with AAV8-RedO-GFP or AAV8-RedO-GFP plus AAV8-RedO-CD47 and PLX5622 from P20-49. Groups without PLX5622 are taken from Figure 2D. Data are shown as mean ± SEM. *** *P*<0.001 by two-tailed Student’s t-test. ns, not significant.

There are two major pathways by which CD47 is known to mediate intercellular signaling. Through its extracellular domain, CD47 can interact with SIRPα, a transmembrane protein expressed on a broad range of cell types including microglia, macrophages, dendritic cells, and neurons (28). In addition, CD47 can be activated by thrombospondin-1 (TSP1), a secreted protein whose binding to CD47 has been shown to help resolve subretinal inflammation (29). To determine if either TSP1-CD47 or CD47-SIRPα signaling were necessary for CD47 to save cones, we generated *rd1*;TSP1^-/-^ and *rd1*;SIRPα^-/-^ mice, which exhibited loss of retinal TSP1 and SIRPα, respectively (Figure 6A). In *rd1*;TSP1^-/-^ animals, co-infection with AAV8-RedO-GFP plus AAV8-RedO-CD47 doubled the number of remaining cones relative to AAV8-RedO-GFP alone (Figure 6, B and C). In contrast, in *rd1*;SIRPα^-/-^ mice, no difference in cone survival was observed with or without AAV8-RedO-CD47. These results demonstrated that the therapeutic effect of AAV8-RedO-CD47 requires SIRPα but not TSP1. Protection of cones with CD47 therefore likely occurs via increased CD47-SIRPα signaling involving one or more non-microglial cell types.

**Figure 6.**
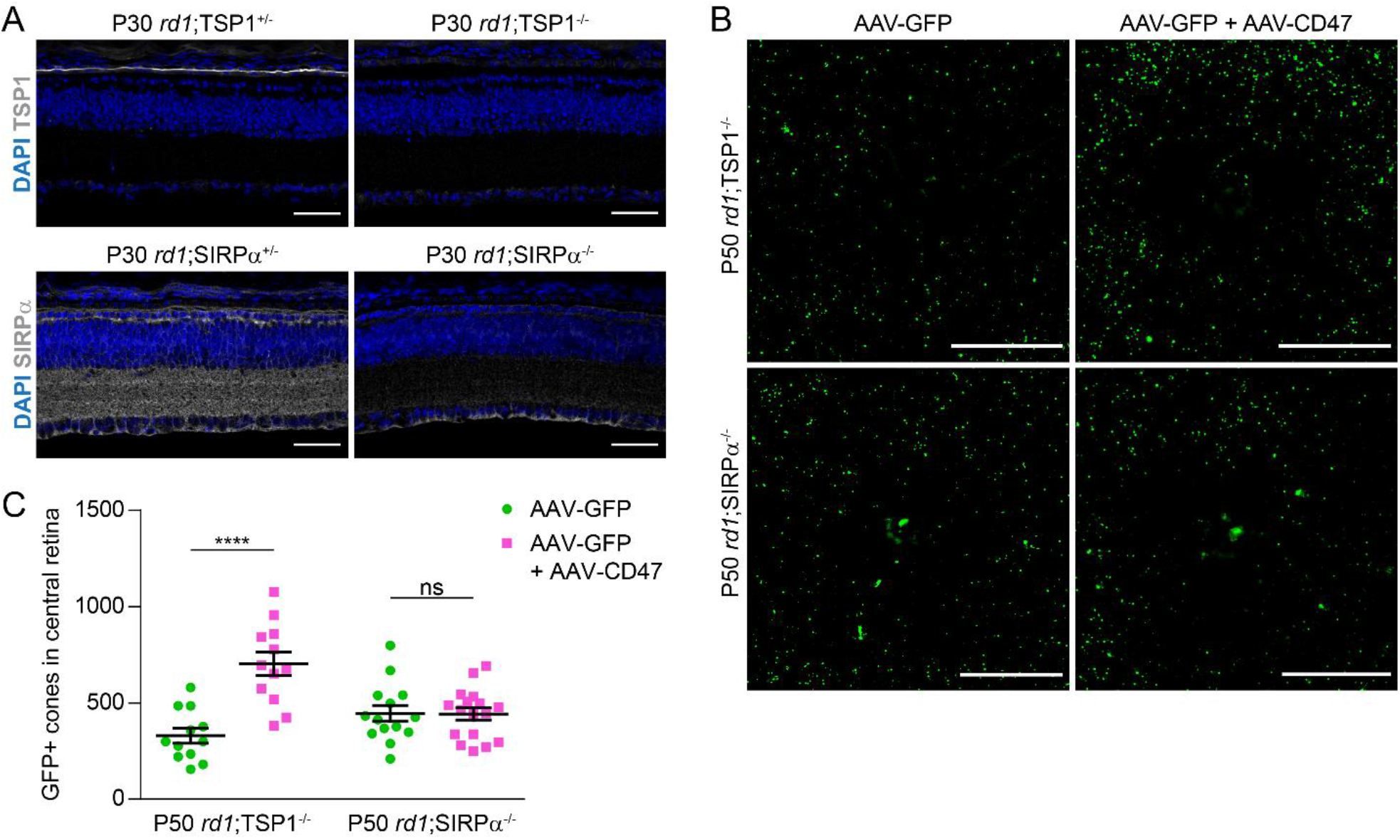
Role of TSP1 and SIRPα in CD47-mediated cone survival. (**A**) Immunostaining for TSP1 and SIRPα, respectively, in P30 *rd1*;TSP1^-/-^ and *rd1*;SIRPα^-/-^ retinas or heterozygous controls. Scale bars, 50 µm. (**B**) Representative images of central retinas from P50 *rd1*;TSP1^-/-^ and *rd1*;SIRPα^-/-^ mice following infection with AAV8-RedO-GFP or AAV8-RedO-GFP plus AAV8-RedO-CD47. Scale bars, 500 µm. (**C**) Quantification of GFP-positive cones in central retinas of *rd1*;TSP1^-/-^ (*n* = 12) and *rd1*;SIRPα^-/-^ (*n* = 14-17) mice following infection with AAV8-RedO-GFP or AAV8-RedO-GFP plus AAV8-RedO-CD47. Data are shown as mean ± SEM. **** *P*<0.0001 by two-tailed Student’s t-test. ns, not significant.

## DISCUSSION

Despite a growing number of gene therapy programs targeting IRDs, the vast majority of patients still lack effective treatment. Correcting individual genes or mutations may improve this situation, but only for subgroups who have those specific lesions, highlighting the need for an intervention that can benefit many more people. In this study, we developed AAV8-RedO-CD47, a novel gene therapy vector that protected cones and vision in multiple models of RP, suggesting it as a potential generic treatment for patients with different mutations. Importantly, delayed expression of CD47 until after most rods had died also preserved cones, implying that this therapy would still be beneficial if administered later in the disease progression. AAV8-RedO-CD47 may thus offer a possible treatment option for many patients with RP, including those who are diagnosed at older ages or with genetics that preclude straightforward gene replacement. The vector might further help combat cone death in other IRDs and degenerative retinal disorders such as age-related macular degeneration (AMD), which affects an estimated 200 million people worldwide (30).

One key question is the mechanism by which CD47 makes cones more resistant to degeneration. While we initially hypothesized that better cone survival would be secondary to reduced engulfment by microglia, this explanation is incompatible with the PLX5622 experiments, in which near complete elimination of retinal microglia failed to perturb CD47-mediated protection of cones. Additionally, although cones may have been exposed to TSP1 secreted from the adjacent retinal pigment epithelium (RPE), AAV8-RedO-CD47 still helped preserve cones in the absence of TSP1-CD47 signaling. Instead, cone rescue with AAV8-RedO-CD47 required SIRPα, indicating that CD47 on cones likely interacts with SIRPα on non-microglial cells to alleviate degeneration. Outside of microglia, SIRPα in the eye is normally expressed in the synapse-rich plexiform layers of the retina (22). However, the protein is also present on multiple immune cell populations, including monocytes, macrophages, neutrophils, and dendritic cells, and has more recently been described on natural killer (NK) and a subset of cytotoxic T cells (28, 31, 32). Binding of CD47 to SIRPα on immune cell populations consistently acts as a negative checkpoint, whether by inhibiting phagocytosis, preventing cell killing, or impeding recruitment of adaptive immunity (28). While Müller glia, another cell type in the retina, have also been shown to phagocytose photoreceptors (33), these cells do not appreciably express SIRPα (34). We therefore suspect that increased CD47-SIRPα signaling with AAV8-RedO-CD47 may save cones by suppressing dysregulated immune cells that would otherwise contribute to cone demise.

Therapeutic interest in the CD47-SIRPα axis has grown tremendously in recent years following the discovery that many cancers co-opt CD47 expression to evade anti-tumor immunity (35). Blocking CD47 and SIRPα have since become promising avenues to induce killing of tumor cells (36, 37), with several modalities now being trialed in patients (38, 39). Rather than disrupt, here we augmented CD47-SIRPα signaling in the eye by using an AAV vector to express CD47. This approach was able to slow progression in multiple models of a blinding disease and may likewise be applicable to other neurodegenerative conditions. For example, CD47 levels have been found to be low in active lesions during multiple sclerosis (40), hinting that increasing CD47 in these regions might ameliorate neuroinflammation. It would similarly be interesting to see if CD47 expression could aid other cell types that undergo degeneration such as retinal ganglion cells in glaucoma or motor neurons in amyotrophic lateral sclerosis (ALS).

## METHODS

### Mice

CD-1 (#022) and FVB (#207) mice were purchased from Charles River Laboratories. CX3CR1^GFP^ (#005582), *rd10* (#004297), C3H (#000659), sighted C3H (#003648), R26-CreERT2 (#008463), and TSP1^-/-^ (#006141) mice were purchased from The Jackson Laboratory. *Rho*^*-/-*^ mice were a gift from Janis Lem (Tufts University) (41). SIRPα^-/-^ mice were a gift from Beth Stevens (Harvard Medical School) (16). CX3CR1^GFP^, R26-CreERT2, TSP1^-/-^, and SIRPα^-/-^ lines were crossed with FVB mice for at least four generations to obtain the following strains: sighted CX3CR1^GFP/+^, *rd1*;CX3CR1^GFP/+^, *rd1*;CreERT2/+, *rd1*;TSP1^+/-^, *rd1*;TSP1^-/-^, *rd1*;SIRPα^+/-^, and *rd1*;SIRPα^-/-^. Genotyping was performed by Transnetyx using real-time PCR. Mice were maintained at Harvard Medical School on a 12-hour alternating light and dark cycle. Animals were housed in standard ventilated racks at a density of up to five per cage. Both males and females were used in all experiments and were randomly assigned to experimental groups.

### Histology

Retinal flat-mounts from *rd1* (FVB), *rd10*, and *Rho*^*-/-*^ mice were prepared as previously described (9, 10). Following dissection of enucleated eyes in phosphate-buffered saline (PBS), retinas were fixed in 4% paraformaldehyde for 30 minutes at room temperature, washed twice with PBS, and relaxed with four radial incisions. For cone arrestin immunostaining, retinas were additionally blocked with 5% donkey serum and 0.3% Triton X-100 in PBS for one hour at room temperature, incubated with 1:3000 of anti-cone arrestin (Millipore) in blocking solution overnight at 4°C, and labeled with 1:1000 of donkey anti-rabbit secondary (Jackson ImmunoResearch) in PBS for two hours at room temperature. Retinas were mounted on microscope slides using Fluoromount-G (SouthernBiotech) with the ganglion cell layer facing up. For retinal cross-sections, enucleated eyes were dissected to remove the cornea, iris, lens, and ciliary body. The remaining eye cups were cryoprotected in a sucrose gradient, frozen in a 1:1 mixture of optimal cutting temperature compound (Tissue-Tek) and 30% sucrose in PBS, and sectioned on a Leica CM3050S cryostat (Leica Microsystems) at a thickness of 20 µm. Tissues were blocked with 5% donkey serum and 0.1% Triton X-100 in PBS for one hour at room temperature and incubated with 1:1000 of anti-CD47 (BD Biosciences), 1:500 of anti-TSP1 (Abcam), or 1:500 of anti-SIRPα (QED Bioscience) in blocking solution overnight at 4°C. Sections were subsequently labeled with 1:1000 of the appropriate secondary antibody (Jackson ImmunoResearch) in PBS for two hours at room temperature followed by 0.5 μg/mL of 4′,6-diamidino-2-phenylindole (DAPI) (ThermoFisher Scientific) in PBS for five minutes at room temperature prior to mounting.

### Ex vivo phagocytosis assay

Freshly dissected retinas were incubated with gentle agitation for one hour at 37°C in 1 mg/mL of pHrodo Red-conjugated zymosan bioparticles (ThermoFisher Scientific) resuspended in retinal culture media (1:1 mixture of DMEM and F-12 supplemented with L-glutamine, B27, N2, and penicillin-streptomycin). Retinas were subsequently washed with PBS and dissociated using cysteine-activated papain as previously described (9). After two additional washes, samples were passed through a 40 μm filter and stained with 0.5 μg/mL of DAPI in FACS buffer (2 mM EDTA and 2% fetal bovine serum in PBS) to exclude non-viable cells. Data were collected on a Cytek DxP11 cytometer and analyzed using FlowJo 10. CX3CR1-positive microglia were defined as phagocytic if positive for pHrodo Red.

### AAV vector cloning and production

The AAV-human red opsin-GFP-WPRE-bGH (AAV8-RedO-GFP) plasmid was a gift from Botond Roska (Institute for Molecular and Clinical Ophthalmology Basel) (42). The AAV8-RedO-CD47 plasmid was cloned by replacing the GFP coding sequence in AAV8-RedO-GFP with that of the most abundant isoform of mouse CD47 (NM_010581.3) (43). The AAV8-RedO-FLEX-mCherry plasmid was cloned by replacing the GFP coding sequence in AAV8-RedO-GFP with that of mCherry inverted and flanked by lox2272 and loxP sites. The AAV8-RedO-FLEX-CD47 plasmid was cloned by replacing the inverted mCherry sequence in AAV8-RedO-FLEX-mCherry with that of inverted CD47. AAV vectors were generated as previously described by transfecting 293T cells with a mixture the vector plasmid, adenovirus helper plasmid, and rep2/cap8 packaging plasmid (44, 45). Seventy-two hours post-transfection, viral particles were harvested from the supernatant, PEGylated overnight, precipitated by centrifugation, treated with Benzonase nuclease (Sigma-Aldrich), and purified through an iodixanol gradient before collection in 100-200 µL of PBS. Vectors were semi-quantitatively titered by SYPRO Ruby (Molecular Probes) for viral capsid proteins (VP1, VP2, and VP3) relative to a reference vector titered by real-time PCR

### AAV vector delivery

Subretinal injections of AAV vectors were performed on neonatal mice (P0-1) as previously described (46). Following anesthetization of the animal on ice, the palpebral fissure was gently opened with a 30-gauge needle and the eye exposed. Using a glass needle controlled by a FemtoJet microinjector (Eppendorf), ∼0.25 μL of vectors were then delivered into the subretinal space. Both left and right eyes were used for injections. AAV8-RedO-GFP was administered at ∼5 × 10^8^ vector genomes (vg) per eye, a dose capable of transducing up to 99% of cones (4). All other vectors were administered at ∼1 × 10^9^ vg per eye.

### Image acquisition and analysis

Retinal flat-mounts were imaged using a Nikon Ti inverted widefield microscope (10x or 20x air objective). Retinal cross-sections were imaged using a Zeiss LSM710 scanning confocal microscope (20x air objective or 40x oil objective). All image analysis was performed using ImageJ. To quantify GFP-positive or cone arrestin-positive cones, custom ImageJ modules were used as previously described (9, 10). For each flat-mount, the locations of the optic nerve head and four retinal leaflets were first manually defined. The number of GFP-positive or cone arrestin-positive objects within the region corresponding to the central retina was then automatically counted and used to represent the number of cones in that sample.

### Light-dark test

Light avoidance in *rd1* (C3H) and sighted C3H mice following no treatment or treatment in both eyes was assessed as previously described (10). Testing was not performed in *rd10* animals as light avoidance in this strain is preserved for at least four months (data not shown). A plastic chamber (Med Associates) measuring 28 cm (length) by 28 cm (width) by 21 cm (height) was divided into two equally sized compartments: one dark and one brightly illuminated (∼900 lux). The two compartments were connected by a small opening and differed in temperature by less than 1°C. At the start of each trial, a mouse was placed in the illuminated compartment and its activity recorded for ten minutes. If after one minute, the animal had not yet entered the dark compartment, it was gently guided there, removed from the chamber, and the trial restarted. Mice were tracked using infrared sensors and location data analyzed with Activity Monitor (Med Associates). Percent time spent in dark was calculated based on the final nine minutes of each trial.

### Optomotor assay

Optomotor responses are absent in the *rd1* strain (47) and were consequently assessed in *rd10* mice. Visual thresholds were measured by an observer (Y.X.) blinded to the treatment groups using the OptoMotry System (CerebralMechanics) as previously described (9). Animals were placed in a chamber with bright background luminance to saturate rods and presented with moving gratings of varying spatial frequencies. Left and right eyes were assessed using clockwise and counterclockwise gratings, respectively, as the optomotor response is evoked by temporal-to-nasal motion in mice (24). For each eye, the highest spatial frequency at which the mouse tracked the grating was determined to be the visual threshold.

### Tamoxifen injections

Tamoxifen (Sigma-Aldrich) was dissolved in corn oil (Sigma-Aldrich) and dosed at 2 mg daily from P19-21 via intraperitoneal injections.

### Microglia depletion

Microglia were depleted using PLX5622 (Plexxikon), an orally available CSF1R inhibitor. PLX5622 was formulated into AIN-76A rodent chow (Research Diets) at 1200 mg/kg and provided *ad libitum* during periods of depletion. For quantification of retinal microglia, freshly dissected retinas were dissociated using cysteine-activated papain as previously described (9). Cells were subsequently blocked with 1:100 of anti-CD16/32 (BD Pharmingen) for five minutes on ice followed by incubation with 1:200 of PE-Cy5-conjugated anti-CD11b (BioLegend), 1:200 of APC-Cy7-conjugated anti-Ly6C (BioLegend), and 1:200 of APC-Cy7-conjugated anti-Ly6G (BioLegend) for 20 minutes on ice. After washes, samples were passed through a 40 μm filter and stained with 0.5 μg/mL of DAPI in FACS buffer. Data were collected on a BD FACSAria II and analyzed using FlowJo 10.

### Statistics

Statistical analyses were performed using GraphPad Prism software. Experimental groups were compared using two-tailed Student’s t-tests, with the addition of a Bonferroni correction if three or more comparisons were performed. A *P* value of less than 0.05 was considered statistically significant. Information on group data and replicates is reported in each figure legend.

### Study approval

All animal experiments were conducted under protocols approved by the Institutional Animal Care and Use Committee (IACUC) of Harvard Medical School.

## AUTHOR CONTRIBUTIONS

S.K.W. and C.L.C. designed the study. S.K.W. and Y.X. performed the experiments and analyzed the data. S.K.W. and C.L.C. wrote the manuscript with input from Y.X.

## ACKNOWLEDGEMENTS

The authors thank Sophia Zhao, the Flow Cytometry Core at Joslin Diabetes Center, the Mouse Behavior Core at Harvard Medical School, and Microscopy Resources on the North Quad at Harvard Medical School for discussions and technical support. This work was supported by the Howard Hughes Medical Institute (C.L.C.) and VitreoRetinal Surgery Foundation (S.K.W.). The authors declare no competing interests.

