## Supplement for "Augmentation of CD47-SIRPα signaling protects cones in genetic models of retinal degeneration"

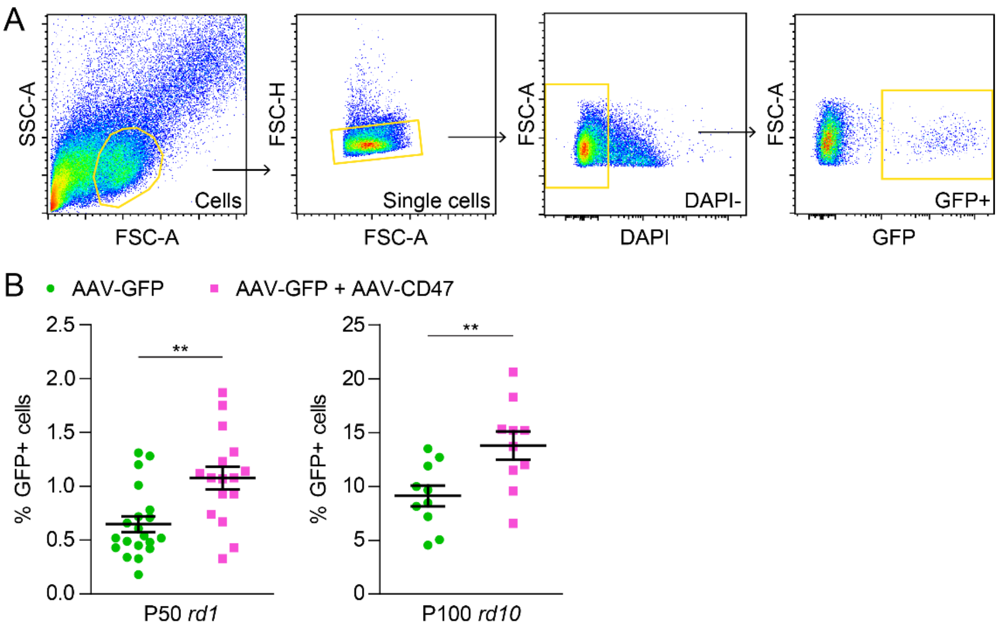


**Supplemental Figure 1. Quantification of cone survival by flow cytometry.**

(**A**) Flow cytometry gating for GFP-positive cones in *rd1* and *rd10* retinas following infection with AAV8-RedO-GFP or AAV8-RedO-GFP plus AAV8-RedO-CD47. (**B**) Quantification by flow cytometry of GFP-positive cones in P50 *rd1* (*n* = 16-20) and P100 *rd10* (*n* = 10) retinas following infection with AAV8-RedO-GFP or AAV8-RedO-GFP plus AAV8-RedO-CD47. Data are shown as mean ± SEM. ** $P<0.01$ by two-tailed Student’s t-test.

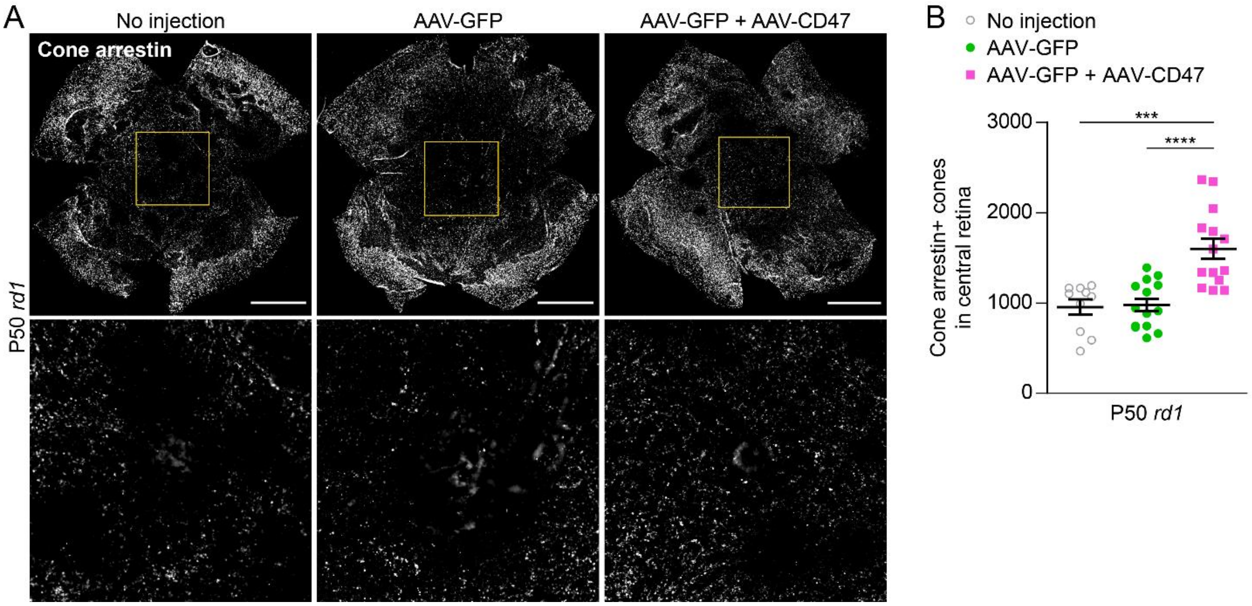


**Supplemental Figure 2. Quantification of cone survival by immunostaining.**

(**A**) Representative flat-mounts of P50 *rd1* retinas without treatment or following infection with AAV8-RedO-GFP or AAV8-RedO-GFP plus AAV8-RedO-CD47 after cone arrestin immunostaining. Paired images depict low and high magnifications. Scale bars, 1 mm. (**B**) Quantification of cone arrestin immunostaining in central retinas of *rd1* mice ($n$ = 10-14) without treatment or following infection with AAV8-RedO-GFP or AAV8-RedO-GFP plus AAV8-RedO-CD47. Data are shown as mean ± SEM. *** $P$<0.001, **** $P$<0.0001 by two-tailed Student's t-test.
